# UNC-16 interacts with LRK-1 and WDFY-3 to regulate the termination of axon growth

**DOI:** 10.1101/2024.02.15.580526

**Authors:** Cody J. Drozd, Tamjid A. Chowdhury, Christopher C. Quinn

**Author notes:** To whom correspondence should be addressed at: Department Biological Sciences, 3209 N. Maryland Avenue, Milwaukee, WI 53211, EMAIL.

## Abstract

*MAPK8IP3 (unc-16/JIP3)* is a neurodevelopmental-disorder associated gene that can regulate the termination of axon growth. However, its role in this process is not well understood. Here, we report that UNC-16 promotes axon termination through a process that includes the LRK-1(LRRK-1/LRRK-2) kinase and the WDFY-3 (WDFY3/Alfy) selective autophagy protein. Genetic analysis suggests that UNC-16 promotes axon termination through an interaction between its RH1 domain and the dynein complex. Loss of *unc-16* function causes accumulation of late endosomes specifically in the distal axon. Moreover, we observe synergistic interactions between loss of *unc-16* function and disruptors of endolysosomal function, indicating that the endolysosomal system promotes axon termination. We also find that the axon termination defects caused by loss of UNC-16 function require the function of a genetic pathway that includes *lrk-1* and *wdfy-3*, two genes that have been implicated in autophagy. These observations suggest a model where UNC-16 promotes axon termination by interacting with the endolysosomal system to regulate a pathway that includes LRK-1 and WDFY-3.

## INTRODUCTION

UNC-16 (JIP3) can serve as a model for understanding how axon transport can regulate axon development and how disruptions in this process can cause neurodevelopmental disorders. UNC-16 is a cytoplasmic adaptor protein that can mediate bidirectional axon transport of vesicles and organelles including mitochondria, golgi, synaptic vesicles, ribosomes, autophagosomes, endosomes and lysosomes (BYRD *et al*. 2001; SAKAMOTO *et al*. 2005; BROWN *et al*. 2009; ARIMOTO *et al*. 2011; EDWARDS *et al*. 2013; EDWARDS *et al*. 2015; NOMA *et al*. 2017; SURE *et al*. 2018; HILL *et al*. 2019; VILELA *et al*. 2019; CELESTINO *et al*. 2022). Moreover, UNC-16 promotes the biogenesis and polarized localization of synapses during axon development (BYRD *et al*. 2001; SAKAMOTO *et al*. 2005; BROWN *et al*. 2009; CHOUDHARY *et al*. 2017). Consistent with its role in neuronal development, rare de novo missense mutations in the human ortholog of UNC-16, JIP3 (also known as MAPK8IP3), are causative for a neurodevelopmental disorder that includes intellectual disability, autism, and developmental delay (IWASAWA *et al*. 2019; PLATZER *et al*. 2019).

The role of UNC-16 (JIP3) in synaptogenesis has been well established. UNC-16 binds to JNK and loss of either protein causes ectopic localization of presynaptic components to dendrites, suggesting that UNC-16 and JNK function together to promote correct localization of presynaptic components (BYRD *et al*. 2001). In addition, other experiments demonstrate that loss of UNC-16 function causes accumulation of RAB-5 and a loss of synaptic vesicles at synapses, suggesting that UNC-16 functions with RAB-5 to promote the biogenesis, or localization of synaptic vesicles (BROWN *et al*. 2009). Consistent with a role in synaptic vesicle biogenesis, UNC-16 can function at the Golgi to sort proteins for inclusion into the synaptic vesicle precursors (CHOUDHARY *et al*. 2017).

In addition to regulating synaptogenesis, UNC-16 can also promote the termination of axon growth (DROZD AND QUINN 2023), though its role in this process is not well understood. Here, we identify genetic interactions between *unc-16, wdfy-3* and *lrk-1* that control axon termination. We report genetic interactions suggesting that UNC-16 functions with the dynein-dynactin complex to promote axon termination. Loss of *unc-16* function causes an accumulation of late endosomes in the distal axon. Moreover, the axon termination defects caused by loss of *unc-16* function are synergistically enhanced by disruptors of the endolysosomal system, including chloroquine, loss of *rab-7* function, and loss of *cup-5* function. We also report that the axon termination defects caused by loss of *unc-16* function can be suppressed by loss of function in a genetic pathway that includes *lrk-1* and *wdfy-3*. Together, these observations suggest a mechanism through which UNC-16 and the endolysosomal system can protect against defects in axon termination. We propose a model whereby UNC-16 prevents axon termination defects through the regulation of LRK-1 and WDFY-3 function.

## RESULTS

### The *unc-16* gene regulates axon termination primarily through its N-terminal short protein isoforms

To investigate the role of UNC-16 in axon targeting, we are using the PLM neuron in *C. elegans*. The PLM cell body resides in the tail and extends an axon along the lateral body wall **(Figure 1A)**. The PLM axon terminates at the midbody, prior to reaching the ALM cell body **(Figure 1A,C)**. Consistent with prior work (DROZD AND QUINN 2023), we have found that a null allele in *unc-16* causes PLM axon termination defects (**Figure 1D-E**), indicating that UNC-16 promotes axon termination. Despite this insight, the mechanistic basis for UNC-16 function in axon termination is not well understood.

**Figure 1:**
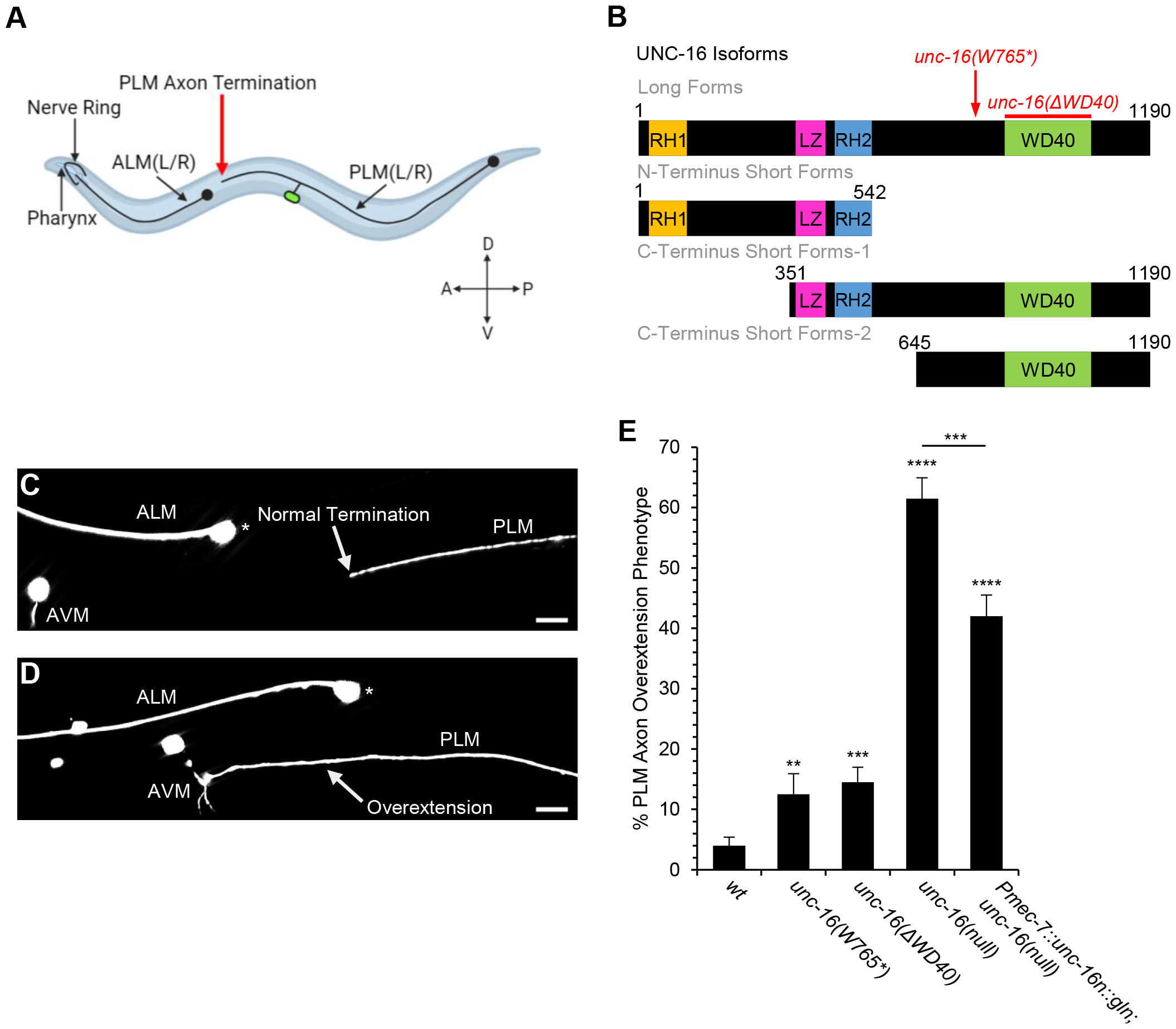
N-terminal short forms of UNC-16 promote axon termination. **(A)** The PLM axon extends a single axon anteriorly that terminates (arrow) prior to the ALM cell body in wild-type *C. elegans*. Illustration created with BioRender.com **(B)** The *unc-16* gene produces 17 protein isoforms (see Supplemental Figure 1). These isoforms can be classified into four groups: the long forms, the N-terminal short forms, the C-terminal short forms-1, and the C-terminal short forms-2. The *unc-16(W765*)* mutation causes a premature stop codon that is predicted to disrupt all UNC-16 isoforms except the N-terminal short forms. The *unc-16(ΔWD40)* mutation is an in-frame deletion of the WD40 domain that is included in all UNC-16 isoforms except the N-terminal short forms. Abbreviations are as follows: RH1 and RH2 = Rab-interacting lysosomal protein (RILP) homology 1 and 2; LZ = leucine zipper. **(C)** An example of normal PLM axon termination in a wild-type PLM, where the axon terminates (arrow) prior to the ALM cell body (asterisk). **(D)** An example of the PLM axon overextension phenotype in an *unc-16* loss-of-function mutant, where the PLM axon terminates (arrow) past the ALM cell body (asterisk). The scale bar represents 10µm. **(E)** The *unc-16(W765*)* and *unc-16(ΔWD40)* mutations cause minimal axon termination defects relative to those caused by an *unc-16(null)* mutation. Introduction of the N-terminus short form partially rescues axon termination defects caused by the *unc-16(null)* mutation. n = 200 axons per phenotype. PLM axons were visualized in L4 stage hermaphrodites with the *jsIs973* transgene that encodes *Pmec-7::rfp*. Asterisks indicate a statistically significant difference, “N-1” Chi-squared test for proportions (**p<0.01, ***p<0.001, ****p<0.0001). Error bars represent the standard error of the proportion. Alleles: *unc-16(W765*)* is unc-16(n730), *unc-16(ΔWD40)* is *unc-16(syb6084)*, and *unc-16(null)* is *unc-16(cue27)*.

To better understand the role of UNC-16 in axon termination, we sought to identify the UNC-16 isoforms that are required for this process. The *unc-16* gene is predicted to give rise to 17 different protein isoforms **(Supplemental Figure 1)**. These isoforms can be grouped into four categories: Long forms, N-terminal short forms, C-terminal short forms-1, and C-terminal short forms-2 **(Figure 1B)**. The long forms include RH1, LZ, RH2, and WD40 domains. The N-terminal short forms include the RH1, LZ, and RH2 domains, but exclude the WD40 domain. The C-terminal short forms-1 include the LZ, RH2 and WD40 domains. The C-terminal short forms-2 include only the WD40 domain.

To determine which UNC-16 forms are important for PLM axon termination, we analyzed *unc-16(n730)* mutants, hereafter called *unc-16(W765*)*. This mutation is expected to remove function of the UNC-16 long forms and both of the C-terminal short forms but spare the N-terminal short forms **(Figure 1B)**. We found that the *unc-16(W765*)* mutation had only a slight effect on axon termination relative to controls **(Figure 1E)**. Since the long forms and C-terminal short forms contain a WD40 domain, we also analyzed mutants that contain an in-frame deletion of the WD40 domain, *unc-16(1ΔWD40)*. We found that this *unc-16(1ΔWD40)* mutation also caused only a slight effect on PLM axon termination **(Figure 1E)**. Since mutations affecting the UNC-16 long form only produced a slight defect, we next asked if an UNC-16 short form is sufficient to promote axon termination. For this experiment, we used the *mec-7* promoter to drive expression in the PLM neuron of UNC-16N::GLN, a C-terminal fusion of mGreenLantern with UNC-16N, one of the six UNC-16 N-terminal short forms. We found that this transgenic expression of UNC-16N::GLN was able to partially rescue the axon termination defects caused by an *unc-16* null mutation, suggesting that the UNC-16 short forms are sufficient to mediate axon termination. Taken together, these observations suggest that axon termination is primarily controlled by the short forms of UNC-16, with a minor contribution from the long forms.

### The dynein-binding function of UNC-16 is required to promote axon termination

Prior work has indicated that a key function of the N-terminal domains of UNC-16 involve binding to kinesin-1 and dynein, both of which are known to bind to the RH1 domain of UNC-16 (CELESTINO *et al*. 2022; SINGH *et al*. 2022). On one hand, kinesin-1 is a motor protein that promotes anterograde transport in axons. On the other hand, the dynein complex functions with the dynactin complex to mediate retrograde transport of vesicles and organelles in axons (RECK-PETERSON *et al*. 2018). Dynein binds to microtubules and functions as a retrograde motor, whereas dynactin binds to dynein and activates its motor activity. Although the interaction between UNC-16 and the dynein is required for normal trafficking of endosomes, the role of this interaction in axon termination has not been investigated.

To investigate a potential role for the dynein-dynactin complex in axon termination, we first asked if dynein or dynactin are required for PLM axon termination. For this experiment, we used a hypomorphic missense mutation in DNC-1 (p150glued), a subunit of dynactin. We found that this *dnc-1* mutation causes PLM axon termination defects **(Figure 2B)**. We also examined a hypomorphic missense mutation in DHC-1 (dynein heavy chain). We found that this DHC-1 mutation also causes PLM axon termination defects. These results suggest that the dynein-dynactin complex is required for axon termination and raise the possibility that the interaction between UNC-16 and dynein is required to promote axon termination.

**Figure 2:**
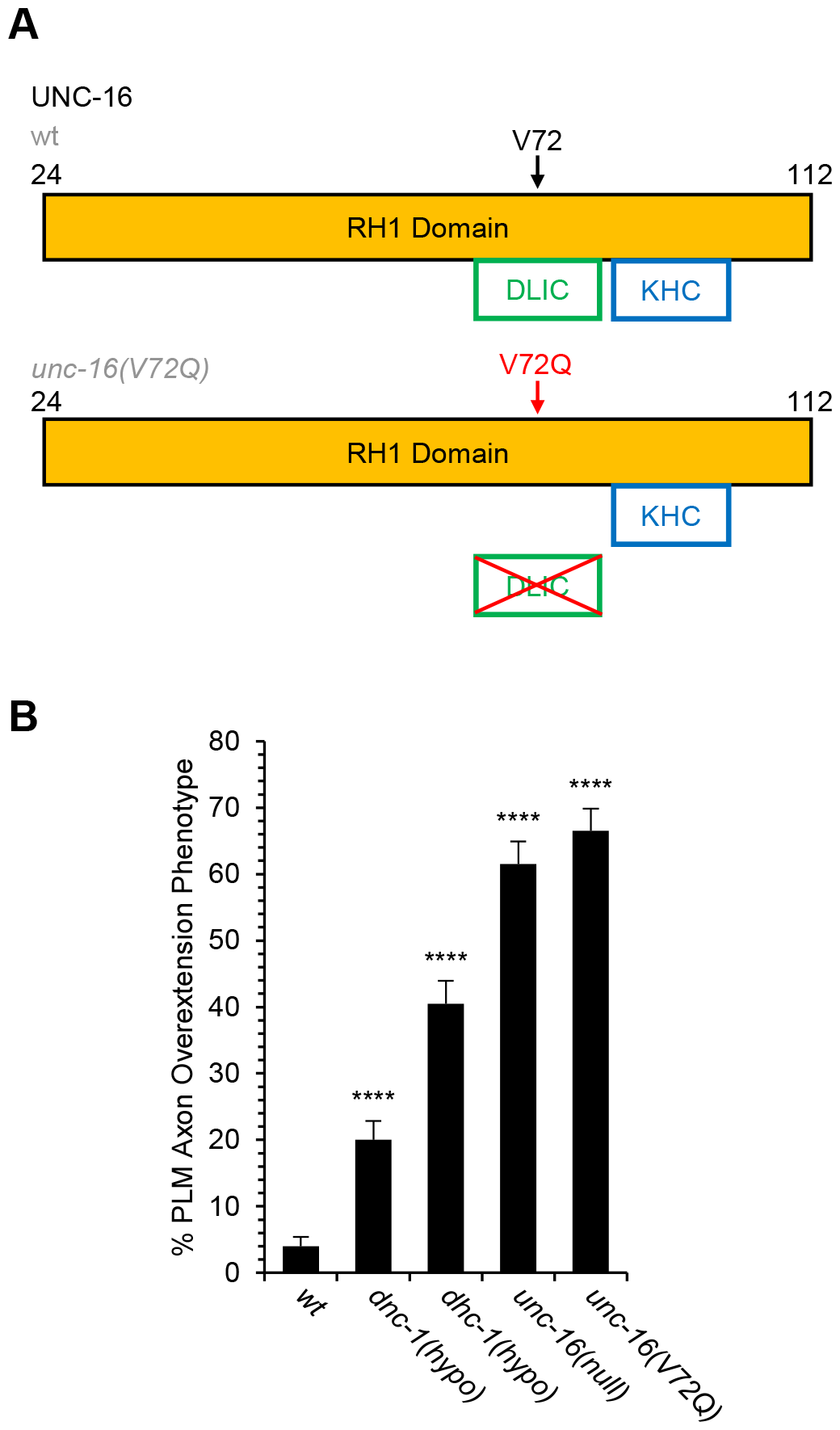
The dynein-binding function of UNC-16 is required to promote PLM axon termination. **(A)** The UNC-16(V72Q) mutation selectively disrupts binding to the dynein light intermediate chain (DLIC). Within the RH1 domain of UNC-16 are binding regions for dynein light intermediate chain (DLIC) and kinesin-1 heavy chain (KHC). In wild-type, DLIC and KHC bind to UNC-16. In the *unc-16(V72Q)* mutant, DLIC is unable to bind but the V72Q mutation does not disrupt the binding between UNC-16 and KHC {Celestino, 2022 #183}. **(B)** The *unc-16(V72Q)* mutation causes PLM axon termination defects equal to those caused by an *unc-16(null)* mutation. Axon termination defects are also caused by a hypomorphic mutation in the DNC-1 ortholog of the p150glued subunit of dynactin and the DHC-1 ortholog of the dynein heavy chain. n = 200 axons per phenotype. PLM axons were visualized in L4 stage hermaphrodites with the *jsIs973* transgene that encodes *Pmec-7::rfp*. Asterisks indicate a statistically significant difference, “N-1” Chi-squared test for proportions (****p<0.0001). Alleles: *dnc-1(hypo)* is *dnc-1(or404), dhc-1(hypo)* is *dhc-1(or283ts)*, and *unc-16(null)* is *unc-16(cue27)*.

To test the idea that UNC-16 interacts with dynein to promote axon termination, we used the *unc-16(V72Q)* mutation. This UNC-16(V72Q) mutant protein selectively disrupts the interaction between UNC-16 and dynein light intermediate chain (DLIC) but leaves the interaction between UNC-16 and the kinesin-1 heavy chain (KHC) intact **(Figure 2A)** (CELESTINO *et al*. 2022). We found that the *unc-16(V72Q)* mutation causes PLM axon termination defects with a penetrance equal to that caused by *unc-16* null alleles **(Figure 2B)**. These observations suggest that the interaction between the UNC-16 RH1 domain and dynein is a key part of the role of UNC-16 in axon termination and further suggest an important role for retrograde transport in this process.

### UNC-16 prevents accumulation of late endosomes in the distal axon

We next characterized the role of UNC-16 in regulating the endolysosomal system within the PLM axon. For these experiments, we focused on late endosomes because we were able to visualize them in wild-type PLM distal axons by using mKate2::RAB-7 as a marker **(Figure 3A-D)** (CELESTINO *et al*. 2022). By contrast, we were unable to visualize early endosomes or lysosomes within the wild-type PLM distal axon. We found that mKate2::RAB-7 puncta are differentially distributed within the wild-type axon, such that their density in the distal axon is about half their density in the proximal axon **(Figure 3A,C,E,F)**. However, loss of *unc-16* function caused accumulation of mKate2::RAB-7 punta in the distal axon, thereby abolishing this differential density of mKate2::RAB-7 puncta **(Figure 3B,D,E,F)**. By contrast, we observed no change in the number of mKate2::RAB-7 in the PLM proximal axon of *unc-16(null)* mutants relative to wildtype **(Figure 3F)**. Taken together, these observations suggest that UNC-16 prevents the accumulation of late endosomes in the distal axon.

**Figure 3:**
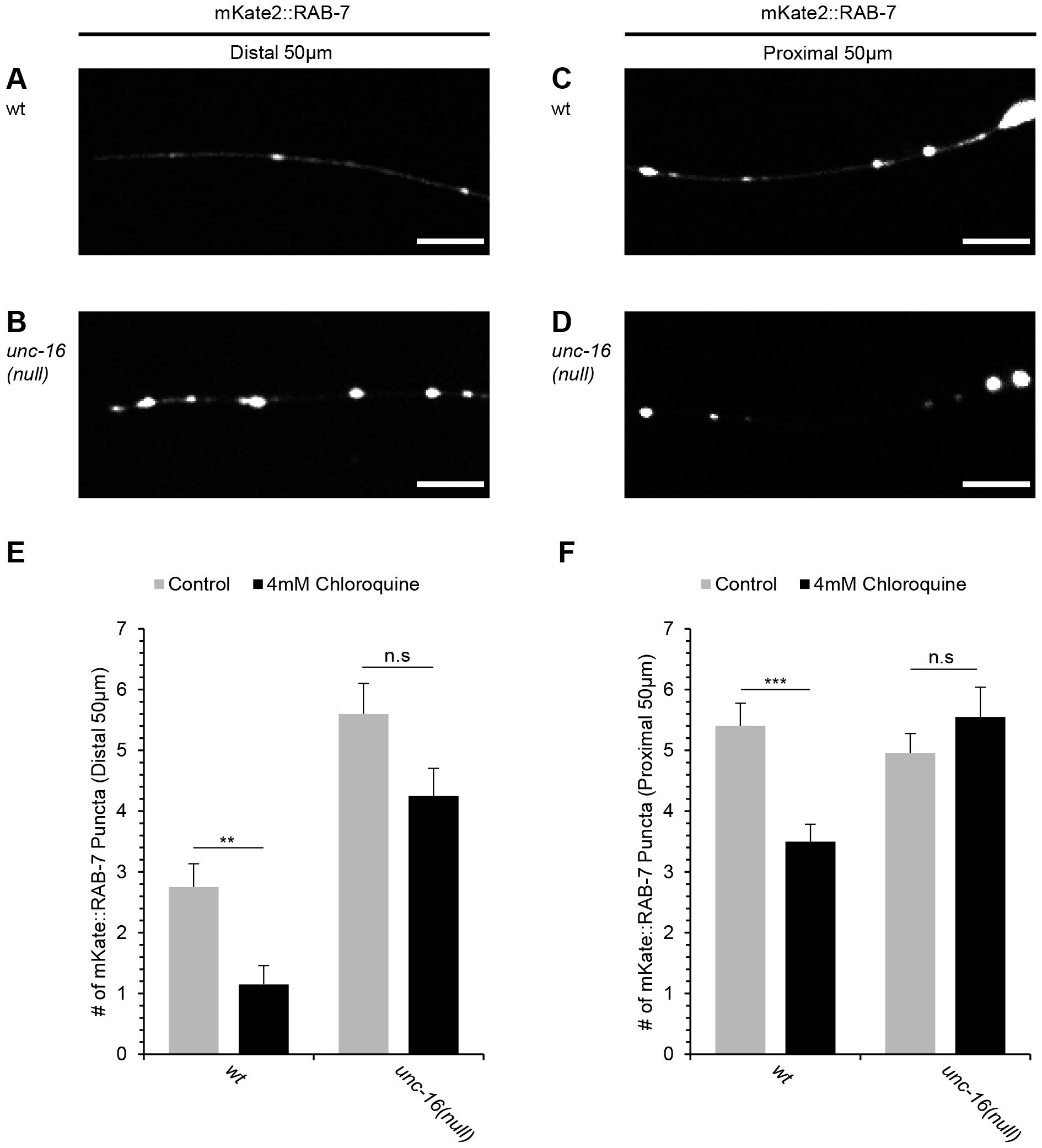
UNC-16-dependant reduction of RAB-7 puncta by chloroquine. **(A)** Example of late endosome localization in a wild-type distal PLM axon. **(B)** Example of late endosome localization in an *unc-16(null)* distal PLM axon. **(C)** Example of late endosome localization in a wild-type proximal PLM axon. **(D)** Example of late endosome localization in an *unc-16(null)* proximal PLM axon. Scale bars represent 10µm. **(E)** The addition of chloroquine reduces the number of RAB-7 puncta in the distal PLM axon but does not affect RAB-7 puncta in the *unc-16(null)* distal PLM axon. **(F)** The addition of chloroquine suppresses the number of RAB-7 puncta in the proximal PLM axon but does not affect RAB-7 puncta in the *unc-16(null)* proximal PLM axon. n=20 axons per genotype. Asterisks indicate a statistically significant difference, independent two-tailed t-test (**p<0.01, ***p<0.001). Error bars represent the standard error of the mean. Late endosomes were visualized with a *mec-7p::mKate::rab-7* transgene. Alleles: *unc-16(null)* is *unc-16(cue27)*.

### UNC-16-dependant reduction of late endosomes by chloroquine

Considering the accumulation of late endosomes in *unc-16(null)* mutants, we hypothesized that UNC-16 promotes the clearance of damaged late endosomes from the distal axon. To test this hypothesis, we used chloroquine, a drug that damages endolysosomal vesicles (MAUTHE *et al*. 2018). We found that treatment with chloroquine, causes a substantial decrease in the number of late endosomes in both the distal and proximal axons of wild-type PLM axons **(Figure 3E,F)**. However, chloroquine caused no significant change in the number of late endosomes in the distal or proximal axons of *unc-16(null)* mutants. These observations indicate that chloroquine reduces the number of mKate2::RAB-7 puncta in an UNC-16-dependant manner, thereby supporting the hypothesis that UNC-16 causes the clearance of damaged late endosomes from the distal axon.

### The endolysosomal system promotes axon termination

Having found that loss of *unc-16* causes accumulation of late endosomes in the distal axon, we next investigated the relationship between endolysosomal function and axon termination. To test for a role of the endolysosomal system in axon termination, we used three different methods to disrupt endosomal function in wild type and *unc-16* mutants. First, we used chloroquine, a lysomyotropic drug that damages lysosomes (MAUTHE *et al*. 2018). We found that 4mM of chloroquine did not cause termination defects in wild type PLM axons **(Fig. 4A)**. However, 4mM of chloroquine did cause a synergistic increase in the penetrance of axon termination defects in *unc-16(null)* mutants. As an alternative method to disrupt endolysosomal system, we also used a maternally rescued null mutation in *rab-7*, which encodes a protein that controls trafficking to and from late endosomes (FENG *et al*. 1995; MUKHOPADHYAY *et al*. 1997). We found that this loss of *rab-7* function did not cause PLM axon termination defects **(Fig. 4B)**. However, loss of *rab-7* function caused synergistic enhancement of the axon termination defects in *unc-16(null)*;*rab-7(lof)* double mutants. Finally, we also disrupted endolysosomal function with a hypomorphic mutation in *cup-5*, a gene that is required for the biogenesis of lysosomes. We found that this *cup-5* mutation did not cause termination defects in wildtype PLM axons, but that it did synergistically enhance defects in *unc-16(null)*;*cup-5(lof)* double mutants **(Fig. 4B)**. These observations suggest that the endolysosomal system promotes axon termination.

**Figure 4:**
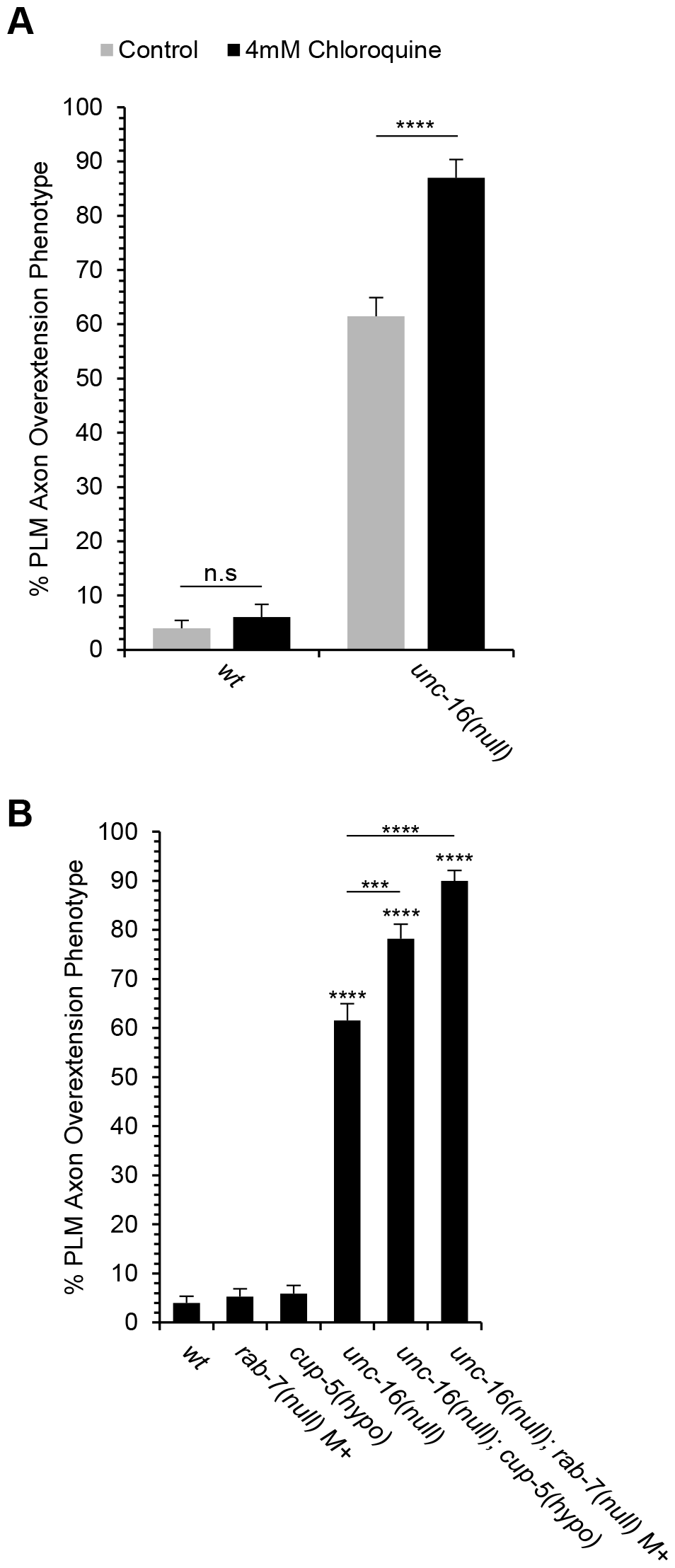
Synergistic interactions between *unc-16* loss of function and disruptors of the endolysosomal system. **(A)** Chloroquine synergistically enhances axon termination defects caused by *unc-16* loss of function. **(B)** Loss of function mutations in *cup-5* and *rab-7* synergistically enhance axon termination defects caused by *unc-16* loss of function. PLM axons were visualized in L4 stage hermaphrodites with the *jsIs973* transgene that encodes *mec-7p::rfp*. Asterisks indicate a statistically significant difference, “N-1” Chi-squared test for proportions (***p<0.001, ****p<0.0001). The *unc-16(null)* and *unc-16(null);rab-7(null) M+* genotypes had an n of 200. The *cup-5(hypo)* and *unc-16(null);cup-5(hypo)* genotypes had an n of 202. The *rab-7(null) M+* genotype had an n of 208. Alleles: *cup-5(hypo)* is *cup-5(ar465), rab-7(null) M+ is rab-7(ok511)* that has been maternally rescued, *unc-16(null)* is *unc-16(cue27)*.

### Axon termination defects caused by loss of UNC-16 function depend on a pathway that includes LRK-1 and WDFY-3

We next used a candidate gene approach to further investigate the mechanism through which UNC-16 protects against neurodevelopmental defects. We initially focused on the LRK-1 endolysosomal protein, which belongs to a family of orthologs including LRRK1/2 in mammals and LrrK in *Drosophila* (hereafter called the LRRK family) *(BISKUP et al. 2006; MONTAGNAC et al. 2009; DODSON et al. 2012)*. In *C. elegans*, LRK-1 has been implicated in axon termination(KUWAHARA *et al*. 2016) and can function with UNC-16 to regulate the trafficking of synaptic vesicle precursors (CHOUDHARY *et al*. 2017). Moreover, the mammalian orthologs of LRK-1 can bind to the mammalian ortholog of UNC-16 (HSU *et al*. 2010).

To test for a genetic interaction between *unc-16* and *lrk-1*, we constructed an *unc-16(null)*;*lrk-1(null)* double null mutant and analyzed PLM axon termination relative to wildtype, *unc-16(null)*; and *lrk-1(null)* single mutants. We found that an *lrk-1(null)* allele caused PLM axon termination defects with a penetrance around 20% **(Figure 5A)**. As mentioned earlier, an *unc-16(null)* allele causes PLM axon termination defects with a penetrance of around 61%. In the *unc-16(null)*;*lrk-1(null)* double mutant, we observed PLM axon termination defects with a penetrance of around 28%, indicating that loss of *lrk-1* function suppresses the axon termination defects that are caused by loss of *unc-16* function. These observations indicate that LRK-1 is required for the axon termination defects caused by loss of *unc-16* function and suggest that UNC-16 can protect against axon termination defects through a mechanism that involves LRK-1.

**Figure 5:**
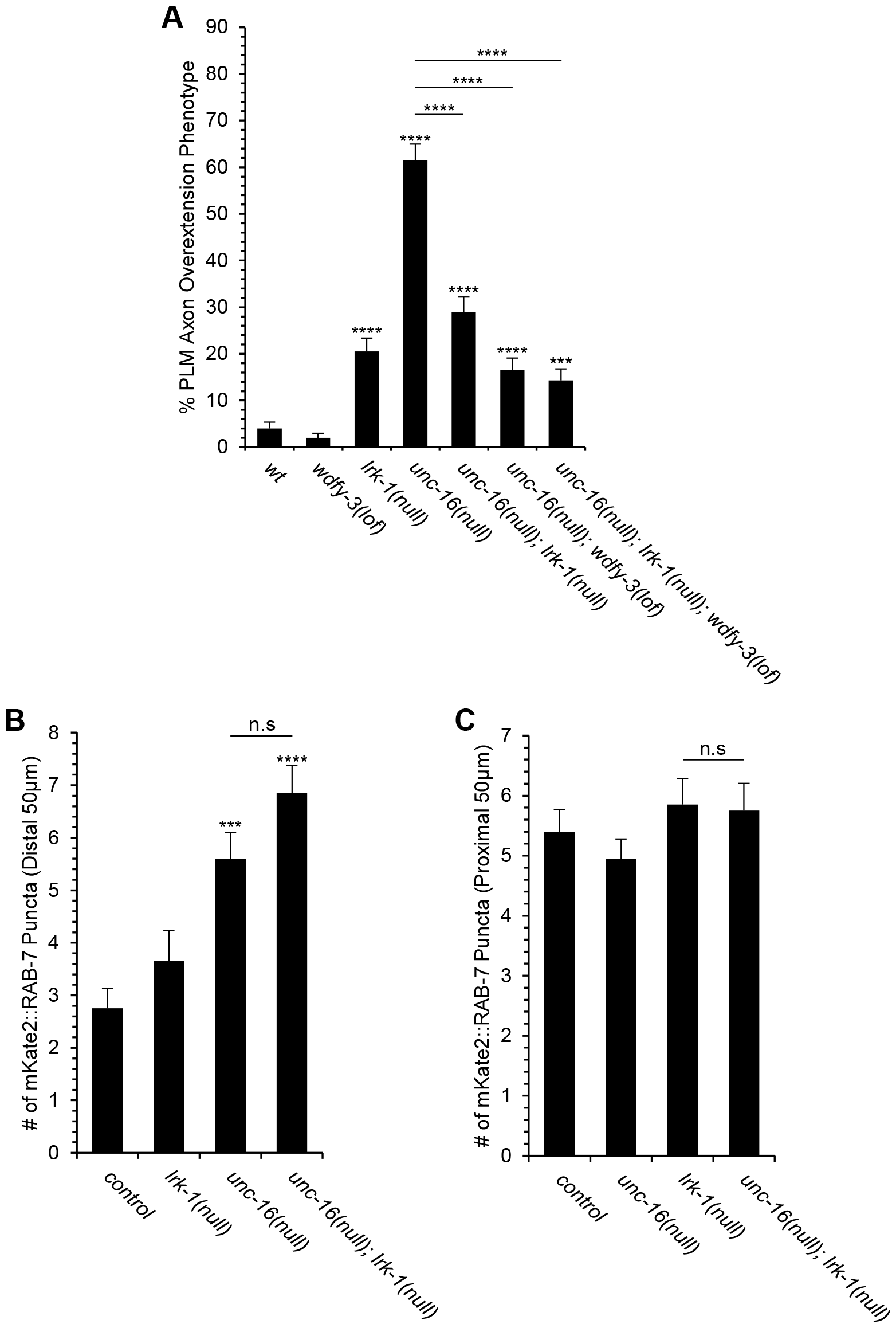
Axon termination defects caused by loss of UNC-16 are dependent on LRK-1 and WDFY-3. **(A)** The axon termination defects caused by loss of *unc-16* function can be suppressed by loss of either *lrk-1* function or loss of *wdfy-3* function. Further suppression does not occur in the unc-16(null);wdfy-3(lof);lrk-1(null) triple mutant. PLM axons were visualized in L4 stage hermaphrodites with the *jsIs973* transgene that encodes *Pmec-7::rfp*. Asterisks indicate a statistically significant difference, “N-1” Chi-squared test for proportions (***p<0.001; ***p<0.0001). The n was 200 for the *unc-16(null)*;*lrk-1(null)*;*wdfy-3(lof)* genotype, 208 for the *wdfy-3(lof)* genotype, and 200 for all other genotypes. **(B)** In the distal axon, loss of *lrk-1* function does not alter the number of late endosomes in either the wild type or *unc-16* null mutant backgrounds. **(C)** In the proximal axon, loss of *lrk-1* function does not alter the number of late endosomes in either the wild type or *unc-16* null mutant backgrounds. Late endosomes were visualized with a *mec-7p::mKate::rab-7* transgene. Asterisks indicate a statistically significant difference, independent two-tailed t-test (***p<0.001, ****p<0.0001). Error bars represent the standard error of the mean. n=20 axons per genotype. Alleles: *lrk-1(null)* is *lrk-1(cue29), unc-16(null)* is *unc-16(cue27)* and *wdfy-3(lof)* is *wdfy-3(ok912)*.

The orthologs of LRK-1 have been implicated in the regulation of late endosomes *(BISKUP et al. 2006; MONTAGNAC et al. 2009; DODSON et al. 2012)*, offering a potential mechanism for the interaction between *unc-16* and *lrk-1* in the control of axon termination. Therefore, we asked if loss of *lrk-1* function alters the number of late endosomes in the distal axon. However, we found no change in the number of late endosomes in the distal or proximal axons of *lrk-1(null)* mutants relative to wild type **(Figure 5B,C)**. Likewise, we also found no change in the number of late endosomes in the distal or proximal axons of *unc-16(null)*;*lrk-1(null)* double mutants relative to *unc-16(null)* single mutants. These observations indicate that loss of *lrk-1* does not suppress *unc-16* axon termination defects by altering the number of late endosomes in the distal axon and suggest that an alternative mechanism is likely to explain the genetic interaction between *unc-16* and *lrk-1*.

Since the LRRK protein family has also been implicated in the regulation of autophagosomes (ALEGRE-ABARRATEGUI *et al*. 2009; BOECKER *et al*. 2021), we considered a role for the WDFY-3 autophagy protein in this process. We focused on WDFY-3, because our previous work indicated that it has a role in the regulation of PLM axon termination (BUDDELL *et al*. 2019). As previously described, we found that loss of *wdfy-3* does not cause PLM axon termination defects **(Fig. 5A)**. However, we found that loss of *wdfy-3* function suppresses the axon termination defects caused by loss of *unc-16* function. These observations indicate that WDFY-3 is required for the axon termination phenotype caused by loss of *unc-16* function.

Having found that mutations in either *lrk-1* or *wdfy-3* can suppress axon termination defects caused by loss of *unc-16* function, we next asked if *lrk-1* and *wdfy-3* function in a genetic pathway to regulate axon termination. To address this question, we constructed a *lrk-1(null)*;*wdfy-3(null)*;*unc-16*(null) triple mutant. We found that the penetrance of PLM axon defects in this triple mutant was no less than either of the double mutants, suggesting that *wdfy-3* and *lrk-1* function in a genetic pathway to regulate axon termination.

### Mutations equivalent to disorder-causing variants in *MAPK8IP3* cause defects in PLM axon termination

Variants in the *MAPK8IP3* ortholog of *unc-16* have been implicated in a rare disorder that is known as the *MAPK8IP3*-related neurodevelopmental disorder, which presents as intellectual disability, developmental delay and autism (BROWN *et al*. 2009; BERGER *et al*. 2017). Moreover, imaging studies have indicated that patients with this disorder also exhibit brain defects including hypoplasia and atrophy of various brain structures (PLATZER *et al*. 2019). The most prevalent defect is in the corpus callosum, which is thinned and shortened relative to healthy controls, suggesting a role for MAPK8IP3 in axonal development in humans.

Considering this role for *MAPK8IP3* in neurodevelopmental disease, we asked if equivalent mutations in *unc-16* would cause defects in PLM axon termination. For these experiments, we screened a panel of five missense mutations in *unc-16* **(Figure 6A)** that are equivalent to variants in *MAPK8IP3* that are associated with the neurodevelopmental disorder (PLATZER *et al*. 2019). We found that in L4 larvae, the UNC-16 L393P mutant protein caused PLM axon termination defects with a penetrance around 19% **(Figure 6B)**. However, none of the other four variants caused PLM axon termination defects in L4 larvae. We also repeated the same screen in L1 larave and found at this stage axon termination could be caused by the L393P mutant protein, the R261Q mutant protein, or by the Y106C mutant protein **(Figure 6C)**. These observations suggest that the neurodevelopmental-disorder associated variants in *MAPK8IP3* can disrupt axon development.

**Figure 6:**
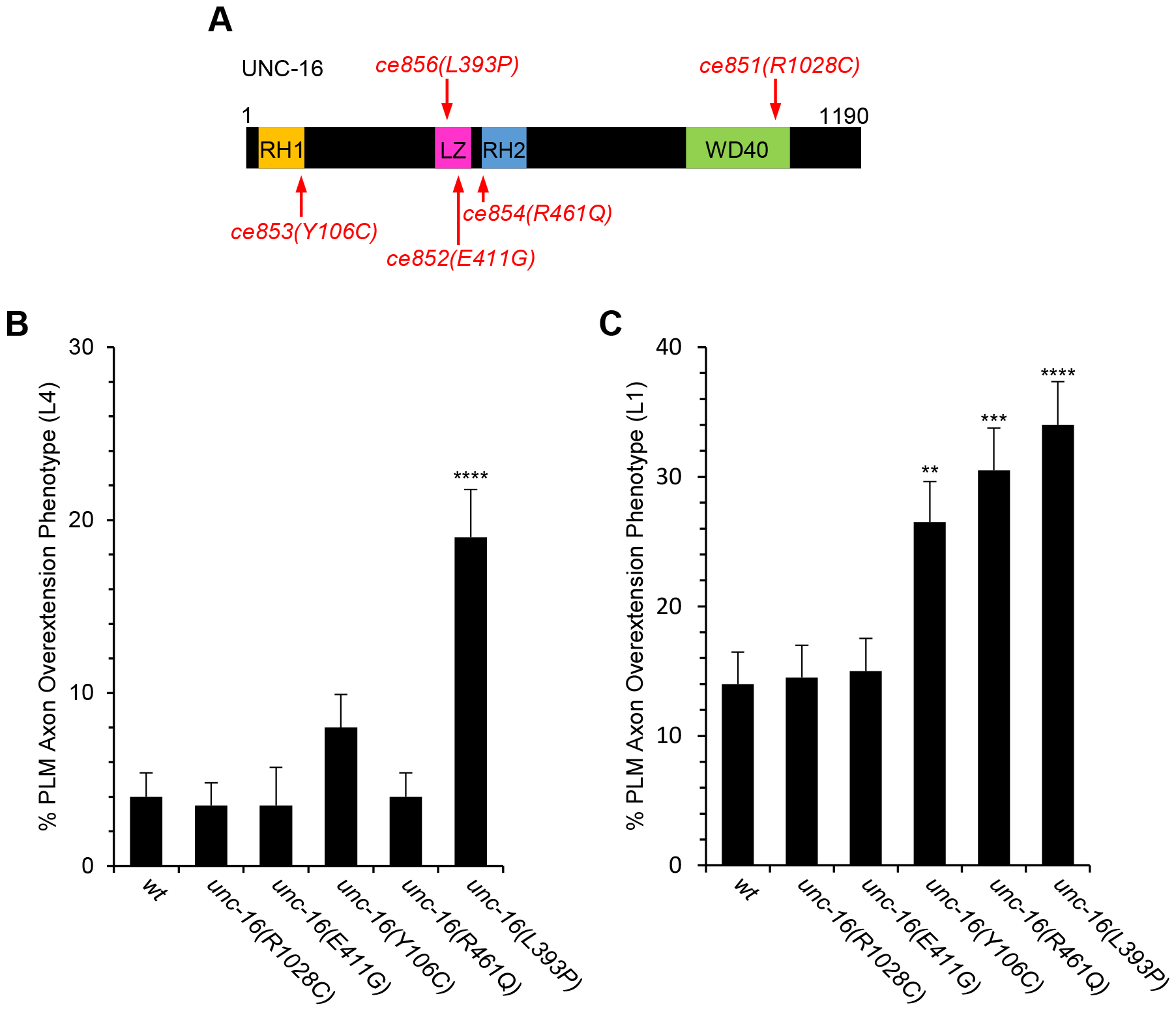
Axon termination is disrupted by *unc-16* mutations that are equivalent to disease-associated variants in *MAPK8IP3*. **(A)** The location of MAPK8IP3-equivilant variants in UNC-16. Abbreviations are: RH1 and RH2 = Rab-interacting lysosomal protein (RILP) homology 1 and 2; LZ = leucine zipper. **(B)** A conserved variant in the LZ domain of UNC-16 causes PLM axon termination defects in the L4 stage. **(C)** Conserved variants in the RH1, RH2, and LZ domains of UNC-16 cause PLM axon termination defects in the L1 stage. PLM axons were visualized in L4 stage hermaphrodites with the *jsIs973* transgene that encodes *Pmec-7::rfp*. Asterisks indicate a statistically significant difference, “N-1” Chi-squared test for proportions (***p<0.0001). n = 200 axons for each genotype. Error bars represent the standard error of the proportion. Alleles: *unc-16(R1028C)* is *unc-16(ce851), unc-16(E411G)* is *unc-16(ce852), unc-16(Y106C)* is *unc-16(ce853), unc-16(R461Q)* is *unc-16(ce854)*, and *unc-16(L393P)* is *unc-16(ce856)*.

## DISCUSSION

Our results support a model for PLM axon termination that involves UNC-16(JIP3), the dynein-dynactin complex, LRK-1(LRRK1/2), and WDFY-3(WDFY3). We report that loss of UNC-16 causes accumulation of abnormally enlarged late endosomes in the distal axon. Moreover, loss of UNC-16 causes PLM axon termination defects that are enhanced by disruptors of the endolysosomal system and suppressed by loss of LRK-1 or WDFY-3, two autophagy-related proteins. Based on these observations, we propose a model whereby UNC-16 promotes clearance of late endosomes from the distal axon. A failure in this process could disrupt axon termination through the misregulation of a pathway that includes LRK-1 and WDFY-3.

### Role of the RH1 domain and the dynein-dynactin complex in axon termination

Our results suggest that the role of UNC-16 in axon termination is mediated primarily by an N-terminal short form of UNC-16, which contains an RH1 domain on its N-terminus and a LZ-RH2 domain combination on its C-terminus. The RH1 domain functions with an adjacent coiled-coil domain to bind and activate the dynein-dynactin complex and this interaction is required for trafficking of endosomes in the axon (CELESTINO *et al*. 2022; SINGH *et al*. 2022). However, the role of this interaction in neuronal development has not been previously addressed.

To address the role of the N-terminal RH1 domain in axon development, we used the *unc-16(V72Q)* mutation to disrupt binding between the RH1 domain and the dynein-dynactin complex. We found that this *unc-16(V72Q)* mutation caused axon termination defects comparable to that caused by *unc-16* null alleles, suggesting that the interaction between the RH1 domain and the dynein-dynactin complex is critical for the role of UNC-16 in axon termination. Consistent with this idea, we also found that hypomorphic mutations in subunits of dynein and dynactin also cause axon termination defects. Together, these observations suggest that the dynein-dynactin complex is required for axon termination and imply that retrograde transport of late endosomes is a key part of this process.

### Role for the endolysosomal system in axon termination

We propose that the role of UNC-16 in axon termination involves the clearance of late endosomes from the distal axon. The role of late endosomes in this process is supported by two observations. First, we found that loss of *unc-16* function causes the accumulation of late endosomes in the distal axon. Second, we observed that disruptors of endosomal function cause synergistic enhancement of axon termination defects caused by *unc-16* loss of function. Therefore, loss of *unc-16* function might compromise the endolysosomal system by altering the trafficking of late endosomes which can be further compromised by chloroquine, loss of *rab-7* function, or loss of *cup-5* function.

The late endosomes that accumulate in the distal axon of *unc-16* mutants are likely to by dysfunctional. This idea is supported by the observation that these late endosomes do not have a normal appearance, but rather are enlarged relative to late endosomes in the distal tip of wild-type axons. Moreover, a role for UNC-16 in clearing damaged late endosomes from the distal axon is supported by our finding that chloroquine reduces the number of late endosomes in wild type axons but not in *unc-16(null)* mutants. Thus, the endosome-damaging drug chloroquine causes a reduction of late endosomes that is dependent on UNC-16 function.

### Role of WDFY-3 and LRK-1 in axon termination

Our results identify a novel genetic pathway that includes *wdfy-3* and *lrk-1*. The WDFY-3 protein is an ortholog of the WDFY3/Alfy protein that has been implicated in autophagy and neuronal development (DRAGICH *et al*. 2016). LRK-1 is an ortholog of LRRK2, which has been extensively studied for its role in Parkinson’s disease and in regulation of the endolysosomal system. In addition, disease-associated mutations in LRRK2 can disrupt autophagy in axons (BOECKER *et al*. 2021). Moreover, emerging data suggest that LRRK2 also has a role in the regulation of neuronal development and de novo mutations in the *LRRK2* gene have been associated with some cases of autism (LABONNE *et al*. 2020; ONISHI *et al*. 2020).

We speculate that the genetic interactions involving *unc-16, wdfy-3*, and *lrk-1* reflect interactions between the endolysosomal system and autophagy. This idea is supported by the fact that the orthologs of *wdfy-3* and and *lrk-1* have both been implicated in the regulation of autophagy and that UNC-16 has been implicated in the regulation of the endolysosomal system (ALEGRE-ABARRATEGUI *et al*. 2009; CLAUSEN *et al*. 2010; BOECKER *et al*. 2021; CELESTINO *et al*. 2022). The endolysosomal system intersects with autophagy, as autophagosomes fuse with late endosomes and lysosomes to create amphisomes and autolysosomes, respectively (TOOZE *et al*. 2014; GANESAN AND CAI 2021). In addition, the endolysosomal system can inhibit autophagy through mTOR localization to lysosomes (YIM AND MIZUSHIMA 2020). Thus, it is possible that disruption of the endolysosomal system by loss of *unc-16* function could cause excess autophagy in the axon. We further speculate that this excess autophagy could lead to defects in axon termination that could be suppressed by loss of function in *wdfy-3* or *lrk-1*.

## Acknowledgements

We thank Reto Gassmann, Kenneth Miller, Michael Nonet, and the Caenorhabditis Genetics Center (funded by NIH P40 OD010440) for strains. We also thank Claire de la Cova for assistance with imaging experiments. This work was funded by the National Institute of Mental Health grant R01MH119157 (to CCQ). This article does not represent the official views of the National Institutes of Health and the authors bear sole responsibility for its content.

## Materials and Methods

### *C. elegans* Genetics and Transgenics

Using standard procedures, *C. elegans* strains were maintained at 20°C on nematode growth medium (NGM)-agar plates. The following alleles were obtained from the CGC: wildtype N2, *unc-16(e109), dnc-1(or404), dhc-1(or283ts), wdfy-3(ok912), unc-16(n730), cup-5(ar465), and rab-7(ok511)*. The *unc-16(n730)* allele is also referred to as *unc-16(W765*)*. The *unc-16(syb6084)* allele is also referred to as *unc-16(ΔWD40)*. This allele was obtained from SunyBiotech and consists of a precise in-frame deletion of the WD40 domain. The *unc-16(cue27) and lrk-1(cue29)* alleles were generated by CRISPR (see below for details). The *unc-16(ce851), unc-16(ce852), unc-16(ce853), unc-16(ce854), unc-16(ce856)* alleles were obtained from Kenneth Miller. Double mutants were constructed using standard methods. Genotypes were confirmed via PCR/sequencing genotypes and associated phenotypes.

The *unc-16(cue27)* allele was constructed using two sgRNAs: one in the 1^st^ intron (5’ – ggccgaggaatagtagtaggggg – 3’) and one in the 3’ UTR (5’ – gatgacaagagaggagacggagg – 3’). Each sgRNA (0.4µg/µL) was added to a mix consisting of 0.8µL Cas9 (6.4µg/µL), 5µL tRNA (0.4µg/µL), and 2µL of an RFP marker. The *lrk-1(cue29)* allele was constructed using two sgRNAs: one in the first exon (5’ – aaaATGGACCTCTCAAGTGGGGG – 3’) and one in the 20^th^ exon (5’ – TCTATTCCATTGCTGTCTGGAGG – 3’). Each sgRNA (0.4µg/µL) was added to a mix consisting of 0.8µL Cas9 (6.4µg/µL), 5µL tRNA (0.4µg/µL), and 2µL of an RFP marker. The *jsIs973* transgene was obtained from Dr. Michael Nonet. The *jsIs973* transgene encodes for *mec-7p::rfp* and was used to examine the PLM axon, ALM cell body, and AVM cell body. The *jsIs821* transgene (obtained from Dr. Michael Nonet) encodes *mec-7p::rab-3::gfp* and was used to examine the clustering of synaptic vesicles at the synaptic branch of the PLM axon. The *Pmec-7::rab-7::mkate2* transgene was obtained from Reto Gassmann and was used to visualize late endosomes.

### Phenotype Analysis

To determine the PLM axon termination phenotype, *C. elegans* were mounted on a 5% agarose pad, anesthetized with levamisole, and observed with a 40x air objective on a Zeiss Axio Imager M2 microscope. A PLM axon was considered overextended if the axon tip terminated anteriorly to the ALM cell body.

To determine the localization of RAB-7::mKate2 in the PLM axon, L4 stage *C. elegans* containing the *mec-7p::rab-7::rfp* transgene were grown at 20°C on NGM-agar plates using standard procedures The number of RAB-7 puncta was analyzed by mounting L4 stage *C. elegans* on a 5% agarose pad, anesthetized with levamisole and observed with a 40x water objective on a Nikon ECLIPSE TI-DH spinning disk confocal microscope. Puncta number was quantified through the use of ImageJ software.

For chloroquine experiments, 100mM chloroquine stock solution was applied to nematode growth medium (NGM)-agar plates and allowed to dry. OP50 was added to dried chloroquine plates. PLM axons were analyzed using the *jsIs973* transgene, which encodes for *mec-7p::rfp*.

### Statistics

Significant PLM axon termination defects were analyzed using the “N-1” Chi-squared test at a 95% CI to compare to the control. 200 axons were analyzed for each allele unless otherwise stated. To determine significant differences among strains with *mec-7p::rab-7::rfp* expression, an unpaired two-tailed t-test with unequal variances was used. 20 axons were analyzed for each allele unless otherwise stated.

**Supplementary Fig 1:**
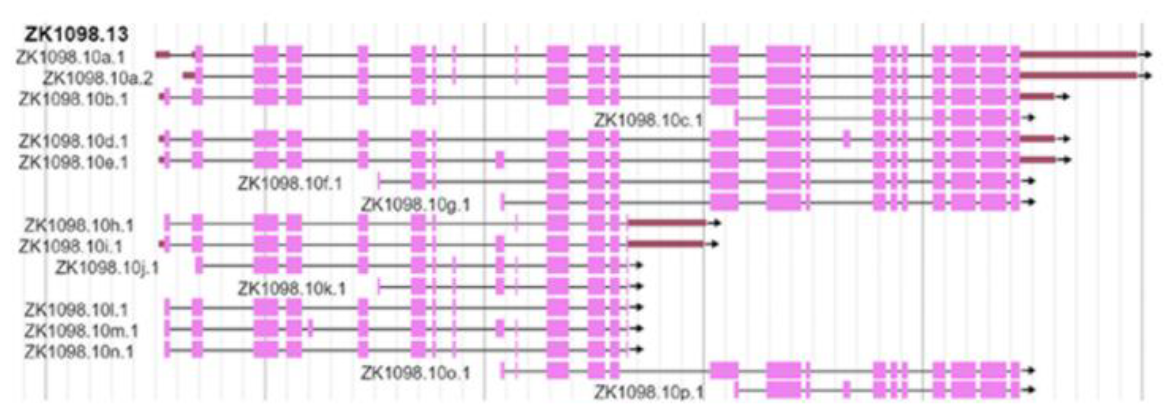
Isoforms of UNC-16. The *unc-16* gene is expressed as 17 protein isoforms. Picture derived from WormBase: https://wormbase.org/species/c_elegans/gene/WBGene00006755#0-9f-10

## Notes

### Competing Interest Statement

The authors have declared no competing interest.

